# Bioinformatics of cyanophycin metabolism genes and characterization of promiscuous isoaspartyl dipeptidases that catalyze the final step of cyanophycin degradation

**DOI:** 10.1101/2023.02.02.526905

**Authors:** Itai Sharon, T. Martin Schmeing

**Affiliations:** Department of Biochemistry and Centre de recherche en biologie structurale, McGill University, Montréal, QC, Canada, H3G 0B1

## Abstract

Cyanophycin is a bacterial biopolymer used for storage of fixed nitrogen. It is composed of a backbone of L-aspartate residues with L-arginines attached to each of their side chains. Cyanophycin is produced by cyanophycin synthetase 1 (CphA1) using Arg, Asp and ATP, and is degraded in two steps. First, cyanophycinase breaks down the backbone peptide bonds, releasing β-Asp-Arg dipeptides. Then, these dipeptides are broken down into free Asp and Arg by enzymes with isoaspartyl dipeptidase activity. Two bacterial enzymes are known to possess promiscuous isoaspartyl dipeptidase activity: isoaspartyl dipeptidase (IadA) and isoaspartyl aminopeptidase (IaaA). We performed a bioinformatic analysis to investigate whether genes for cyanophycin metabolism enzymes cluster together or are spread around the microbial genomes. Many genomes showed incomplete contingents of known cyanophycin metabolizing genes. Cyanophycin synthetase and cyanophycinase are usually clustered together when recognizable genes for each are found within a genome. Cyanophycinase and isoaspartyl dipeptidase genes typically cluster within genomes lacking *cphA1*. About one-third of genomes with genes for CphA1, cyanophycinase and IaaA show these genes clustered together, while the proportion is around one-sixth for CphA1, cyanophycinase and IadA. We used X-ray crystallography and biochemical studies to characterize an IadA and an IaaA from two such clusters. The enzymes retained their promiscuous nature, showing that being associated with cyanophycin-related genes did not make them specific for β-Asp-Arg dipeptides derived from cyanophycin degradation.

## Introduction

Cyanophycin is a biopolymer first described over 100 years ago as large, light scattering granules observed in cyanobacterial cells[1]. These granules are composed of chains with backbones of L-aspartate residues with L-arginine attached to each Asp side chain[2] (Fig. 1**a**). Cyanophycin contains 26% nitrogen content by mass, which, along with its inert nature and low solubility, makes it useful for nitrogen, carbon and energy storage[3–5]. Cyanophycin can be produced by a wide variety of bacteria[6, 7], but research in a biological context has mostly focused on cyanobacteria[8–12]. Cyanophycin is known to be especially useful for nitrogen-fixing cyanobacteria, which separate anaerobic nitrogen fixing from oxygen-producing photosynthesis either spatially in different cell types[8] or temporally in a day/night cycle[13].

**Figure 1.**
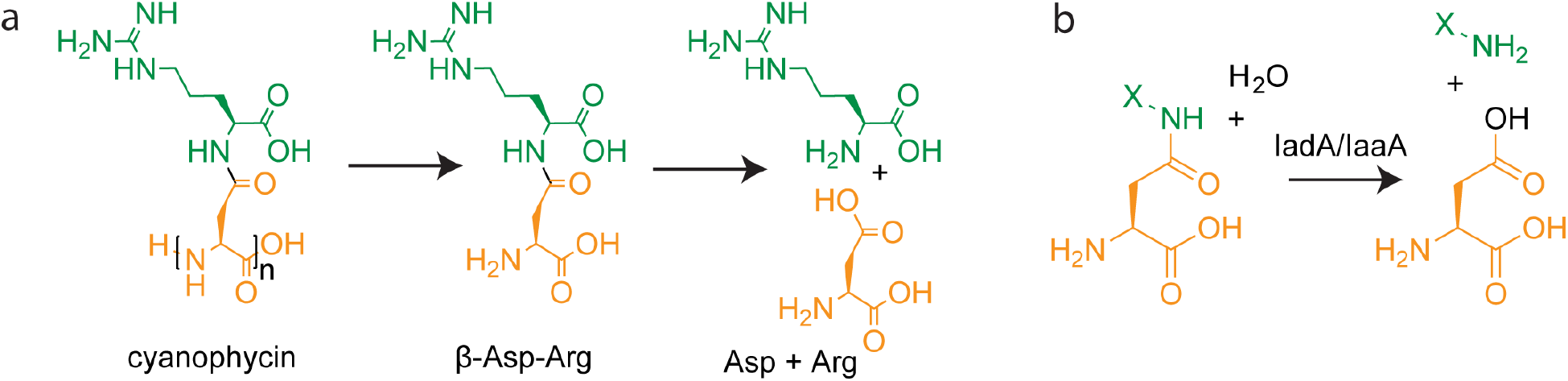
The structure and degradation of cyanophycin. (**a**) Long polymer chains (typically n=80-400) are degraded by cyanophycinase into β-Asp-Arg dipeptides, which are then hydrolyzed by isoaspartyl dipeptidases, resulting in free Asp and Arg. (**b**) The general reaction is catalyzed by isoaspartyl dipeptidases. X=any amino acid.

Cyanophycin is made by cyanophycin synthetase 1 (CphA1)[14] or 2 (CphA2)[15] (supplementary Fig. S1). CphA1 is a widespread enzyme that catalyzes two ATP-dependent reactions[14, 16]: it first adds Asp to the polymer backbone and then attaches Arg to the side chain of that Asp residue through an isopeptide bond[6]. Some CphA1 enzymes can also incorporate lysine into cyanophycin in place of arginine, though at lower efficiency[17]. CphA2, a cyanobacterial enzyme related to CphA1, uses a single active site to catalyze the ATP-dependent repolymerization of β-Asp-Arg dipeptides into cyanophycin[15, 18].

To access the nitrogen, carbon and energy stored in cyanophycin[8, 13], bacteria degrade it into free amino acids. This is done in two steps (Fig. 1**a**, supplementary Fig. S1): First, cyanophycin is hydrolyzed into β-Asp-Arg dipeptides by a specialized exo-cyanophycinase enzyme, either the intracellular CphB[19] or CphI[7], or the extracellular CphE[19]. Then the β-Asp-Arg dipeptides are hydrolyzed into Asp and Arg by enzymes that possess isoaspartyl-dipeptidase activity[20] (Fig. 1**b**). The two degradation steps occur within the same cells in cyanobacterial species that have day/night regulation of cyanophycin metabolism[21], while in cyanobacterial communities with cyanophycin-synthesizing heterocysts, dipeptides are shuttled to vegetative cells for hydrolysis[8]. Many bacterial communities capable of using exogenous cyanophycin as a carbon and nitrogen source have been identified[22, 23]. These communities can be found in a variety of environments, such as animal gut flora[24], soil[25] and fresh-water sediments[26], suggesting cyanophycin is commonly found in these environments. There is evidence that the two steps of cyanophycin degradation are sometimes split between members of a bacterial consortium, where some members express cyanophycinase and others degrade the β-aspartyl dipeptides[22].

Enzymes capable of degrading β-aspartyl dipeptides are very common, because β-aspartyl residues can form spontaneously from intramolecular rearrangement of Asp and Asn residues in proteins[27]. The resulting β-aspartyl dipeptides, if not degraded, can accumulate to pathological levels in cells[28]. In bacteria, these β-aspartyl residues can either be repaired by L-isoaspartyl O-methyltransferase enzymes (E.C 2.1.1.77)[29] or be hydrolyzed into their amino acid constituents[30]. Two bacterial enzymes are known to have significant β-aspartyl dipeptidase activity: isoaspartyl dipeptidase (IadA)[31, 32], a bacterial zinc metallopeptidase; and isoaspartyl aminopeptidase (IaaA, also called plant-type asparaginase, EcAIII and IadC)[20, 33–35], a common Ntn-family enzyme with known plant and animal homologs. IadA and IaaA are evolutionarily unrelated and have different catalytic mechanisms, but both have broad substrate specificity because damage to proteins can lead to the attachment of different amino acids to Asp/Asn side chains[20, 32, 36]. Accordingly, they are also capable of degrading β-Asp-Arg/Lys, so it is assumed that β-Asp-Arg/Lys dipeptides derived from cyanophycin are degraded by general isoaspartyl dipeptidases[7, 19, 20, 37]. In addition, several other enzymes, such as glycosylasparaginases, catalyze similar reactions and can display low levels of β-aspartyl dipeptidase activity[38].

In this study, we analyzed the genomes in the NCBI RefSeq database[39] to investigate the tendency of cyanophycin metabolism genes to co-occur and cluster together in the genome. We observe moderate levels of co-occurrence of cphA1, cyanophycinase and an isoaspartyl dipeptidase genes within these genomes. The rates of clustering of various combinations of the genes are well above random, ranging from moderate (e.g., 37 of 231 genomes containing *cphA1*, a cyanophycinase gene and *iadA* show all three genes to cluster) to high (e.g., 30 of 32 genomes with a cyanophycinase gene and *iaaA*, but without *cphA1* genes show clustering). Characterization of the activity and structures of representative IadA and IaaA enzymes which cluster with cyanophycin synthetase and cyanophycinase genes revealed that they have not become specific for β-Asp-Arg dipeptides.

## Results

### Identification of cyanophycin-metabolizing gene clusters

To quantify the occurrence and clustering tendency of cyanophycin-metabolizing genes, we analyzed the presence and genomic localization of *cphA1*, cyanophycinase (*cphB, cphI* or *cphE*) and isoaspartyl dipeptidase (*iaaA[20]* or *iadA*[31]) in all 27,349 non-redundant, complete bacterial genomes in the NCBI RefSeq database[39]. Isoaspartyl dipeptidases are common (found in 11,814 genomes, 43.2%), which is expected, as they have roles other than cyanophycin metabolism. Cyanophycin synthetase 1 is found in 1,614 genomes (6%, Table 1), and a recognizable cyanophycinase gene is present in 739 genomes (3%, Table 2).

**Table 1.** Analysis of genomes which encode CphA1.

|  | CphA1 +<br>cyanophycinase<br>and/or laaA<br>and/or ladA | CphA1 +<br>cyanophycinase | CphA1<br>+ laaA | CphA1<br>+ ladA | CphA1 +<br>laaA<br>and/or<br>ladA | CphA1 + laaA +<br>cyanophycinase | CphA1 + ladA +<br>cyanophycinase | CphA1 +<br>cyanophycinase<br>+ laaA and/or<br>ladA |
| --- | --- | --- | --- | --- | --- | --- | --- | --- |
| In genome | 1614 | 658 | 968 | 232 | 1181 | 153 | 231 | 366 |
| Clustered | 538 | 535 | 51 | 38 | 88 | 49 | 37 | 85 |

**Table 2.** Analysis of genomes which encode a cyanophycinase.

|  | cyanophycinase<br>+ CphA1 and/or<br>laaA and/or ladA | cyanophycinase<br>+ laaA | cyanophycinase<br>+ ladA | cyanophycinase<br>+ laaA and/or<br>ladA |
| --- | --- | --- | --- | --- |
| In genome | 739 | 185 | 251 | 418 |
| Clustered | 578 | 79 | 52 | 130 |

Next, we examined the tendency of cyanophycin-metabolizing genes to co-occur and cluster together. We defined co-occurrence as at two genes present in the same genome, and clustering as genes separated by not more than a 5 kilobase pair (kbp) intergenic region. Of the genomes that have *cphA1*, 658 (41%) also have a recognizable cyanophycinase. These genes are clustered in most (535; 82%) of the genomes that have both. Genes for IaaA or IadA are found in 1181 (73%) *cphA1*-containing genomes, with 968 (60%) of those genomes having *iaaA* and 232 (14%) having *iadA*. However, in contrast to cyanophycinase genes, isoaspartyl dipeptidase genes generally do not cluster with *cphA1*, being proximal in only 88 (7.5%) of genomes that have both (Table 1).

Interestingly, clustering of *cphA1* and isoaspartyl dipeptidase is more common in genomes that have genes encoding all three steps of cyanophycin metabolism. There are 366 such genomes in the RefSeq database. In genomes that have *cphA1*, a cyanophycin gene, and *iaaA*, 49 of 153 show clustering. In the case of *iadA*, 37 of 231 genomes with *cphA1*, a cyanophycin gene and *iadA* show these three clustered.

Ben Hania et al. have described the utililty and occurance of a “cyanophycin utilization locus” which includes cyanophycinase genes, *iadA* and a transporter so a microbe can scavenge cyanophycin from the environment[40]. This observation also holds for *iaaA*: Searches of the NCBI RefSeq database returned 52 genomes that contain a cyanophycinase gene and *iaaA* or *iadA* but not *cphA1*, and 45 of them had cyanophycinase and isoaspartyl-dipeptidase genes clustered (Table 3).

**Table 3.** Analysis of genomes which encode a cyanophycinase and isoaspartyl dipeptidase but not CphA1.

|  | cyanophycinase<br>+ laaA (no<br>cphA) | cyanophycinase<br>+ ladA (no<br>cphA) | cyanophycinase<br>+ laaA and/or<br>ladA (no cphA) |
| --- | --- | --- | --- |
| In genome | 32 | 20 | 52 |
| Clustered | 30 | 15 | 45 |

The rate of each of the above clusterings is above random chance: As a control, we detected 955 genomes with *cphA1* and dihydrofolate reductase (*folA*), a common housekeeping gene unrelated to cyanophycin metabolism. None of these genomes had the two genes clustered together.

### IadA and IaaA from cyanophycin clusters are not specific for β-Asp-Arg/Lys

Previous studies which characterized the activity of canonical isoaspartyl dipeptidases found that both IadA[32] and IaaA[20] accept a wide range of β-aspartyl dipeptides as substrates. Subsequent structural results explained this lack of substrate specificity: while both enzymes make extensive interactions with the Asp portion of the substrate, the portion of the isoaspartyl dipeptidase surrounding the amino acid attached to the Asp side chain is large and able to accommodate the substrate rather than bind it specifically[32, 41].

We wondered whether the IaaA or IadA homologs present in cyanophycin metabolism clusters have evolved to specialize in cyanophycin degradation and display substrate preference for β-Asp-Arg (and β-Asp-Lys) over other β-aspartyl dipeptides. We therefore performed biochemical and structural characterization of a representative of IaaA and of IadA β-aspartyl dipeptidases whose genes are clustered with both *cphA1* and *cphB*: IadA from *Leucothrix mucor* DSM2157 (*Lm*IadA) and IaaA from *Roseivivax halodurans* DSM15395 (*Rh*IaaA).

*Lm*IadA has 44% sequence identity to *E. coli* IadA (*Ec*IadA[32]). Like *Ec*IadA, the purified enzyme forms octamers in solution (supplementary Fig. S2)[32]. We examined the activity of *Lm*IadA towards several β-aspartyl dipeptides and found that it displayed no apparent preference towards β-Asp-Arg/Lys (Fig. 2**a**). To confirm the structural basis for this lack of specificity, we solved the structure of the wild type enzyme at 1.8 Å resolution and compared it to that of *Ec*IadA[32] (Table S1).

**Figure 2.**
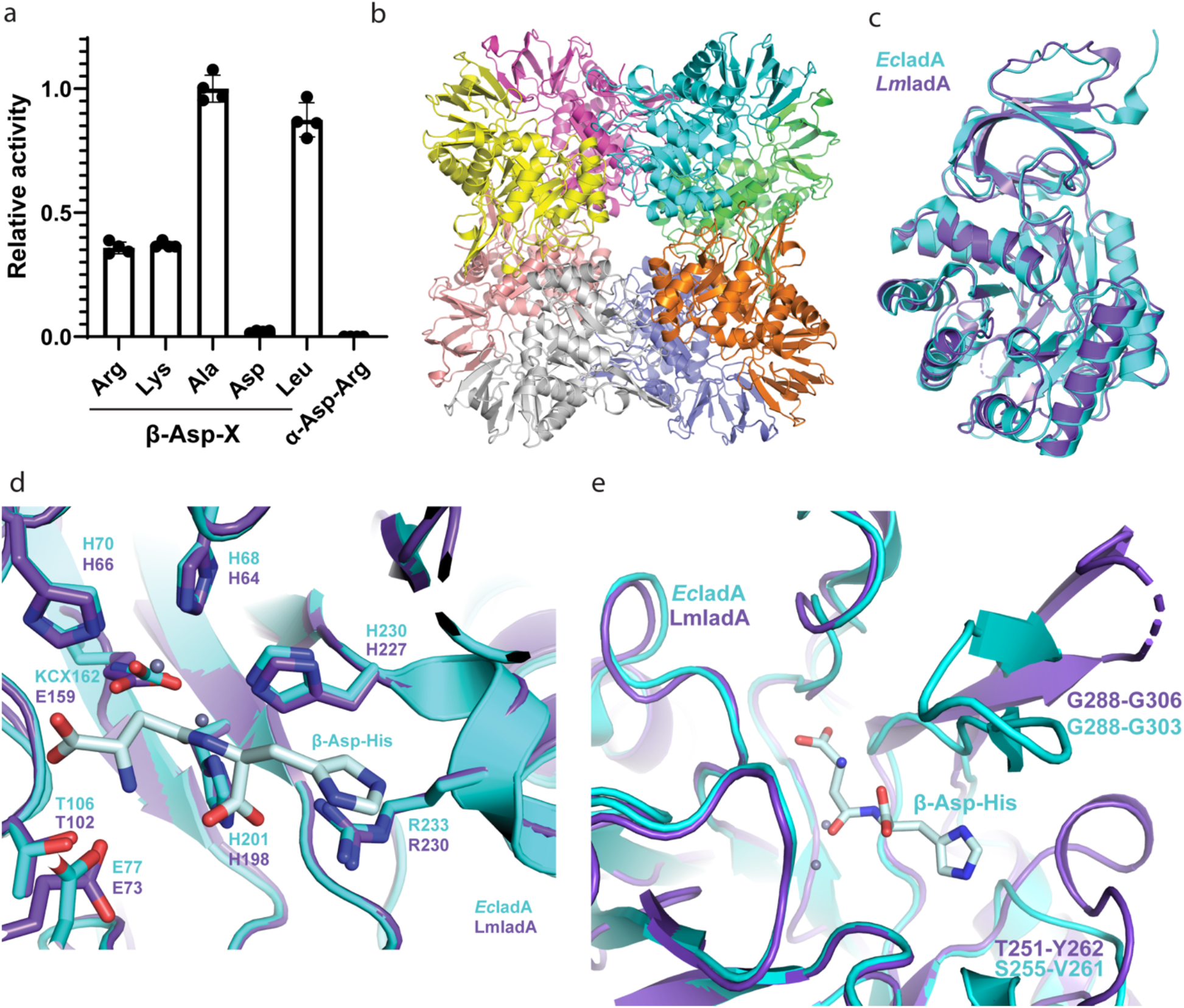
Structure and activity of *Lm*IadA. (**a**) Asp release assay of *Lm*IadA and different Asp-containing dipeptides. The enzyme is specific towards β-aspartyl dipeptides, but displays no specificity towards Arg or Lys as the β-linked amino acid. Error bars represent the standard deviation of the mean of n=4 replicates. (**b**) The homooctameric crystal structure of *Lm*IadA. (**c**) Overlay of *Lm*IadA (purple) and *Ec*IadA[32] (cyan, PDB code 1YBQ) monomers showing their high overall structural similarity. (**d**) Close-up view of the active sites of *Lm*IadA and *Ec*IadA in complex with the substrate β-Asp-His, showing they are similar in both sequence and structure. (**e**) Overlay of the regions around the active sites of *Lm*IadA and *Ec*IadA, showing both have large openings capable of accommodating a variety of β-aspartyl dipeptides as substrate.

The crystal structure of *Lm*IadA shows a homooctameric architecture as the asymmetric unit (Fig. 2**b**). It displays high similarity to that of *Ec*IadA[32] (0.81 Å RMSD across 315 Cα pairs, PDB code 1YBQ; Fig. 2**c**), with the active site residues almost identical in both sequence and structure (Fig. 2**d**). Two Zn^2+^ ions are liganded by H64, H66, H198, H227 and E159, corresponding to *Ec*IadA H68, H70, H201, H230 and carboxylated K162. Substrate binding residues in *Ec*IadA such as E77, T106 and R233[32], are also present at corresponding positions in *Lm*IadA (E73, T102 and R230) and display similar conformations (Fig. 2**d**).

The published structure of *Ec*IadA in complex with β-Asp-His[32] shows that the His side chain of the substrate forms minimal interactions with the enzyme. It faces an opening in the active site which, as expected, can accommodate a variety of substrates. *Lm*IadA displays a somewhat different architecture in this region (Fig. 2**e**). The loop formed by *Lm*IadA T251-Y262 is longer and bulkier than the corresponding one of *Ec*IadA (S255-V261), and as a result could restrict access to the active site. However, the partially flexible region between G288-G306 (*Ec*IadA G288-G303) is oriented away from the binding pocket. This leads to a similarly sized opening in the active site region surrounding the non-Asp portion of the substrate and explains the lack of specificity (Fig. 2**e**).

We performed analogous analyses with the IaaA enzyme encoded in the cyanophycin gene cluster of *Roseivivax halodurans. Rh*IaaA has 51% sequence identity with *E. coli* IaaA (*Ec*IaaA[41]). Like *Ec*IaaA and other Ntn-family enzymes, the pro-enzyme is expressed as a single chain that undergoes autocatalytic cleavage into two subunits, a and b, which constitute the mature a2b2 heterotetrameric enzyme (supplementary Fig. S2). We assayed the activity of *Rh*IaaA towards the same set of β-Asp dipeptides used to assess *LmIadA* and found that it could hydrolyze all of them with no apparent preference towards β-Asp-Arg/Lys (Fig. 3**a**). To confirm the structural basis for the lack of substrate specificity, we solved the structure of the wildtype enzyme at 2.7 Å resolution and compared it to that of *Ec*IaaA (supplementary Table S1).

**Figure 3.**
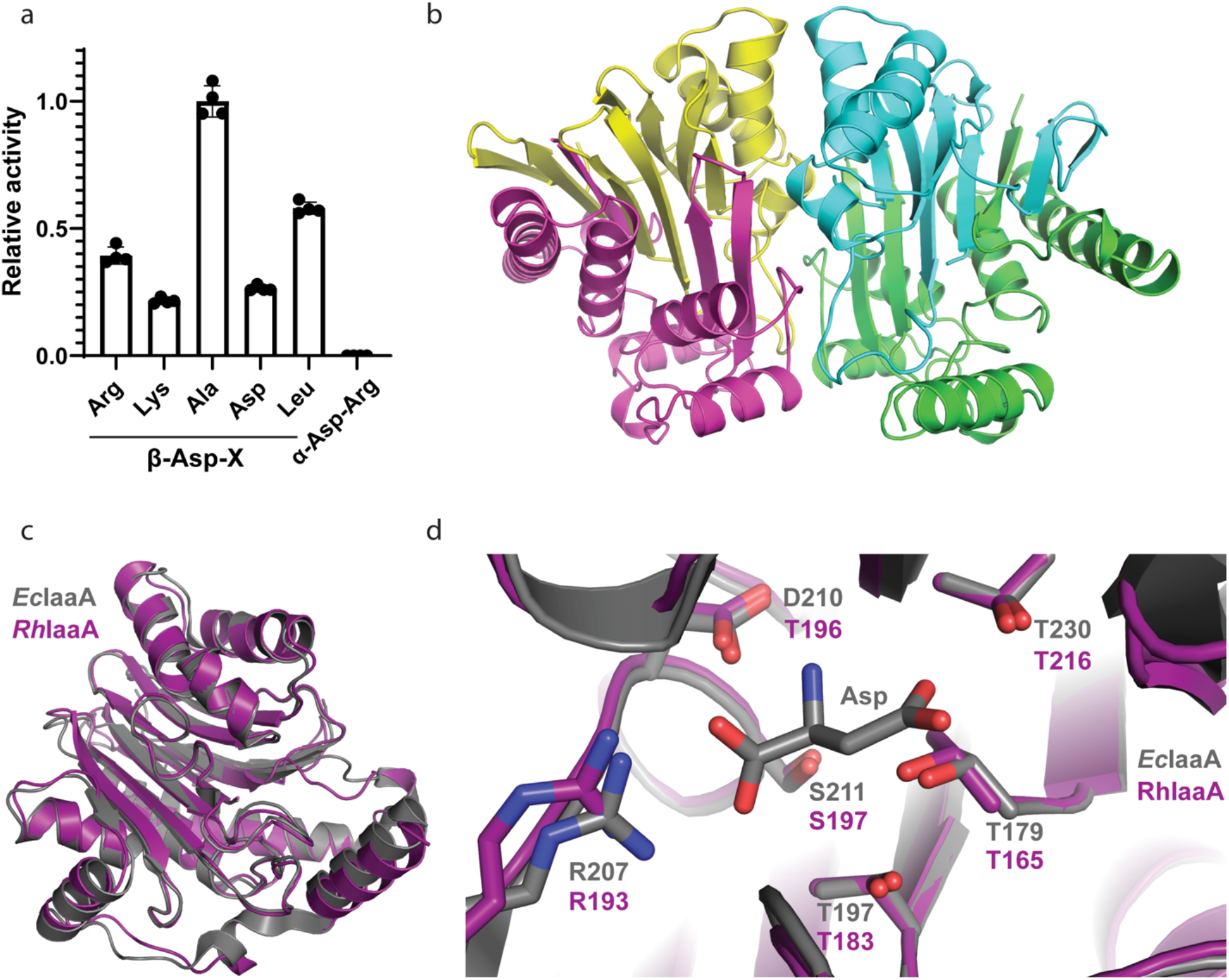
Structure and activity of *RhIaaA*. (**a**) Asp release assay of *RhIaaA* with different Asp-containing dipeptides. The enzyme is specific towards β-aspartyl dipeptides, but displays no specificity towards Arg or Lys as the β-linked amino acid. Error bars represent the standard deviation of the mean of n=4 replicates. (**b**) The heterotetrameric crystal structure of *Rh*IaaA. (**c**) Overlay of *Rh*IaaA (purple) and *Ec*IaaA[41] (gray, PDB code 2ZAL) heterodimers showing their high overall structural similarity. (**d**) Close view of the active sites of *Rh*IaaA and *EcIaaA* in complex with the product Asp, showing they are similar in both sequence and structure.

The crystal structure of *Rh*IaaA shows the expected heterotetrameric architecture (Fig. 3**b**). The enzyme displays high structural similarity to *Ec*IaaA (0.58 Å RMSD across 230 Cα pairs, PDB code 2ZAL[41]; Fig. 3**c**), with the active site residues being almost identical in both sequence and conformation. In *Ec*IaaA, the Asp portion of the substrate is bound by T197, R207, D210, S211, T230 and G231, as well as the catalytic T179[41]. These residues are all present and in the same conformations in *Rh*IaaA (T183, R193, D196, S197, T216 and G217, and the catalytic T165, Fig. 3**d**). As is the case with IadA, the substrate likely binds oriented in a way that positions the non-Asp portion of it facing a large opening in the active site (Fig. 3**d**). This presumably results in minimal interaction between IaaA and the substrate residue bound to Asp by the scissile isopeptide bond, which would enable the active site to accommodate a wide range of β-aspartyl dipeptides.

## Discussion

Bacteria often use clustering to control expression of genes with related functions[42]. In the case of cyanophycin metabolism, clustering appears to be common for *cphA1* and cyanophycinase[9] (Table 1). Previous studies in cyanobacteria show that these two genes can also share some transcription regulation elements[9]. Clustering of genes for cyanophycinase and an isoaspartyl dipeptidase is very common in genomes that have those genes but not *cphA1* (Table 2). These are often accompanied by amino acid transporters and probably represent cyanophycin-scavenging clusters, such as the ones described in the cyanobacteria-scavenger strain L21-Spi-D4[40] and in *Flammeovirga pacifica* strain WPAGA1[43].

The clustering rate of isoaspartyl dipeptidases with *cphA1* and cyanophycinase in genomes that have all three is well above random distribution, but not as high as that of *cphA1*-cyanophycinase alone. There are several possible explanations why clustering is not strict. First, it is possible for these genes to be under control of the same transcription regulators even if they are not clustered. Second, since isoaspartyl dipeptidases are required outside of a cyanophycin context, there may be evolutionary pressure to keep those genes separate for regulatory purposes. Third, in some cases it is beneficial to have cyanophycin-metabolizing genes regulated independently of one another. An example for this can be seen in the heterocyst-forming cyanobacterium *Anabaena* sp. PCC7120. Heterocysts of this bacterium express cyanophycinase to degrade cyanophycin into dipeptides, which are shuttled to vegetative cells. These, in turn, express high levels of IaaA to convert the dipeptides into free amino acids[8].

In general, the co-occurrence rates of genes involved in cyanophycin metabolism is lower than we expected. For example, detection of a recognizable cyanophycinase in only 41% of *cphA1*-containing genomes is unanticipated. Cyanophycin is only known to serve as a storage material, and the bacteria that store it must also possess the wherewithal to degrade it. It is possible that bacteria which possess *cphA1* but not *cphB/E/I* possess other, unknown cyanophycinase isozymes. The lack of an identifiable isoaspartyl dipeptidase gene in 27% of *cphA1*-containing genomes suggests that not all genes encoding enzymes with this dipeptidase activity were detected in our searches. Similarly, Füser *et al*. performed an analysis of 48 *cphA1* or cyanophycinase-containing genomes in 2007[7] and found that only 26 also had *iaaA* or *iadA*. Isoaspartyl dipeptidase activity in these bacteria could be provided by distant homologues of *iaaA* or *iadA* or by unrelated isozymes. Manual examination of genomes from the RefSeq database that have a CphA1-cyanophycinase cluster shows some of them to include adjacent genes which could potentially have isoaspartyl dipeptidase activity, such as those annotated as “S9 family peptidase” (in genome NZ_CP029187.1), annotated as “M14 family metallopeptidase” or “succinylglutamate desuccinylase/aspartoacylase family protein” (in genome NZ_VYQF01000002.1) and a gene weakly homologous (25-30% identity) to cocaine esterase[44] (in genome NZ_SJEY01000003). The existence of cryptic isoaspartyl dipeptidase enzymes has been proposed before, for example in *Saccharomyces cerevisiae*[45].

Both of the isoaspartyl dipeptidases from cyanophycin gene clusters that we cloned, expressed, purified and assayed display no substrate specificity towards β-Asp-Arg/Lys and accept a range of isoaspartyl dipeptides. The crystal structures of both enzymes were consistent with this promiscuity and show that the structural basis for this lack of specificity is shared with other IaaA and IadA enzymes. These results suggest that even when their genes are clustered with cyanophycin-related genes, IaaA and IadA function in both cyanophycin metabolism and the protein-degradation pathway, in line with the widely held belief that general isoaspartyl dipeptidases are responsible for the last step of cyanophycin degradation[7, 20].

## Methods

### Bioinformatics

For the identification of gene clusters, we created a local database with all complete bacterial genomes in the NCBI (USA) Refseq[39] database (May 2022). We used cblaster[46] to search this database using several queries for *cphA1* (WP_028947105.1, WP_004925893.1, WP_015942562.1), cyanophycinase (WP_011058003.1, WP_004925892.1, Q8KQN8.1), *iadA* (WP_188415469.1, WP_138978951.1, WP_037265155.1) and *iaaA* (MBS3792760.1, WP_034545427.1, WP_022952024.1). For the identification of putative isoaspartyl dipeptidases in *cphA1*-cyanophycinase clusters, MultiGeneBlast[47] was used to search *cphA1*-containing genomes for *cphA1*-cyanophycinase clusters, and the results were analyzed manually for putative isoaspartyl dipeptidases.

### Cloning, protein expression and purification

The genes encoding *Lm*IadA (WP_022952024.1) and *Rh*IaaA (WP_037265155.1) were amplified from genomic DNA (DSMZ, Leibniz Institute, Germany). Both genes were cloned into a plasmid derived from pJ411 with a a C-terminal tobacco etch virus (TEV) protease cleavage site and a 8xHis affinity tag. Gene subcloning and mutagenesis were performed by transforming PCR fragments with overlapping ends into chemically competent DH5-α *E*. coli cells. Proteins were expressed in *E. coli* BL21(DE3) cells grown in TB media supplemented with 150 μg/ml kanamycin. Cultures were grown at 37 °C until they reached an OD_600_ of ~1. The growth temperature was then lowered to 18 °C and protein expression was induced with 0.2 mM isopropyl β-d-1-thiogalactopyranoside (IPTG) for ~20 hours. All subsequent protein purification steps were carried out at 4 °C. Following harvest by centrifugation, the cells were resuspended in buffer A (250 mM NaCl, 50 mM Tris pH 8.0, 10 mM imidazole, 2 mM β-mercaptoethanol) supplemented with a few crystals of lysozyme and DNase I, and lysed by sonication. The lysate was clarified by centrifugation at 40,000 g for 30 minutes and then applied onto a HisTrap HP column (Cytiva, USA). The column was washed extensively with buffer B (buffer A with 30 mM imidazole) and the protein was eluted with buffer C (buffer A with 250 mM imidazole). For structural studies, the protein was incubated with TEV protease for removal of the octahistidine tag while being dialyzed overnight against buffer D (250 mM NaCl, 20 mM Tris pH 8.0, 5 mM β-mercaptoethanol) prior to application to a HisTrap column and collection of the flow through. All protein preparations were then concentrated and applied to a Superdex200 16/60 column (Cytiva, USA) equilibrated in buffer E (100 mM NaCl, 20 mM Tris pH 8.0, 1 mM dithiothreitol). Fractions with the highest protein purity were concentrated, supplemented with glycerol to a final volume of 15% and flash frozen in liquid nitrogen for storage.

### Protein crystallization, data collection, structure solution and refinement

For crystallization trials, all proteins were buffer exchanged into buffer E and subjected to small-scale wide screen crystallization trials in 96-well plates using the sitting drop method. Optimization of crystallization conditions was performed using the sitting drop method by mixing 2 μl of protein with 2 μl of crystallization buffer and allowing this to equilibrate against 500 μl of crystallization buffer. The crystallization buffer for *Lm*IadA (20 mg/ml) contained 0.56 M NaH2PO4 and 1.04 M K2HPO4. Crystals were grown at 22 °C and cryo-protected by briefly dipping them in crystallization solution supplemented with 20% glycerol before freezing in liquid nitrogen. Data were collected at the Advanced Light Source (ALS) beamline 5.0.1. The structure was solved by molecular replacement using *E. coli* IadA (PDB code 1YBQ) as a search model. The crystallization buffer for *Rh*IaaA (10 mg/ml) contained 0.1 M bis-tris propane pH 8.5, 0.2 M disodium malonate and 25% PEG3350. Crystals were grown at 4 °C and cryo-protected by dipping them in crystallization solution supplemented with 10% PEG100 for 1 minute before freezing in liquid nitrogen. Data were collected at the Canadian Light Source (CLS) beamline CMCF-BM. The structure was solved by molecular replacement using *E. coli* IaaA (PDB code 2ZAL) as a search model. All datasets were processed in DIALS[48] and merged in AIMLESS[49] implemented in CCP4i2 suite[50]. The structures were refined in REFMAC5[51], Rosetta[52], Phenix[53] and Coot[54]. Figures were prepared in PyMOL (Schrödinger, USA).

### Enzyme activity assays

Enzyme-catalyzed β-Asp-X dipeptide hydrolysis was measured with an Asp release assay[32]. The 100 μl reactions contained 100 mM HEPES pH 8.2, 20 mM KCl, 5 mM α-ketoglutarate, 1 mM NADH, 2.4 U aspartate aminotransferase, 0.3 U malate dehydrogenase, 1 mM dipeptide substrate and 500 nM purified enzyme. Data were collected by following 340 nm transmittance in 96-well plates using a SpectraMax Paradigm (Molecular Devices, USA) and analyzed using Prism (GraphPad, USA). β-Asp-Arg dipeptides were purified as previously described[18]. β-Asp-Ala and α-Asp-Arg were purchased from Bachem (Switzerland). β-Asp-Lys and β-Asp-Leu were purchased from Toronto Research Chemicals (Canada). β-Asp-Asp was purchased from Advanced ChemBlocks (USA).

## Acknowledgements

We thank all the members of the Schmeing lab for important advice and ongoing discussions on this project, Nancy Rogerson for proofreading and synchrotron staff J. Gorin (Canadian Light Source) and M. Allaire (Advanced Light Source) for facilitating remote collection of diffraction datasets. Part of the research described in this paper was performed using beamline CMCF-BM at the Canadian Light Source, a national research facility of the University of Saskatchewan, which is supported by the Canada Foundation for Innovation (CFI), the Natural Sciences and Engineering Research Council (NSERC), the National Research Council (NRC), the Canadian Institutes of Health Research (CIHR), the Government of Saskatchewan, and the University of Saskatchewan. Beamline 5.0.1 of the Advanced Light Source, a U.S. DOE Office of Science User Facility under Contract No. DE-AC02-05CH11231, is supported in part by the ALS-ENABLE program funded by the National Institutes of Health, National Institute of General Medical Sciences, grant P30 GM124169-01. This work was funded by CIHR Project Grant 178084 and a Canada Research Chair to TMS.

## Data Availability

Diffraction data and structures determined in this study have been deposited to the Protein Data Bank: *Lm*IadA (PDB 8DQN; https://www.rcsb.org/structure/8DQN), *Rh*IaaA (PDB 8DQM https://www.rcsb.org/structure/8DQM). All other relevant data are within the manuscript and its Supplementary Information files.

## Author contribution

I.S. and T.M.S. designed the study and performed the bioinformatic analysis. I.S. performed all biochemical and structural experiments and data processing. I.S. and T.MS. wrote the manuscript.

## Additional information

The authors declare no competing interests.

