## Supplemental Material for "Bioinformatics of cyanophycin metabolism genes and characterization of promiscuous isoaspartyl dipeptidases that catalyze the final step of cyanophycin degradation"

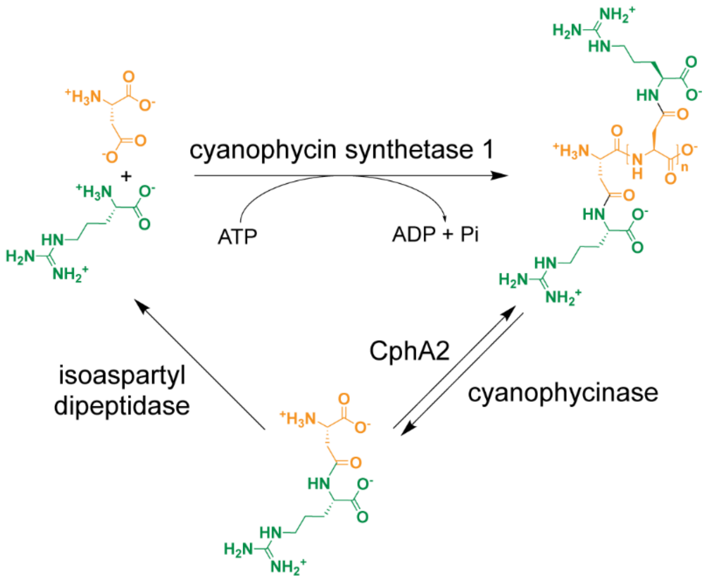


**Figure S1.** Schematic diagram of cyanophycin biosynthesis and degradation.

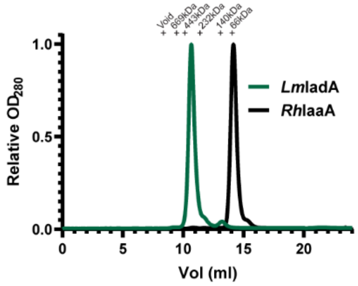


**Figure S2.** Size exclusion chromatography traces of *Lm*IadA and *Rh*IaaA suggesting they migrate as octamer (expected Mw 335 kDa) and heterotetramer (expected Mw 66 kDa), respectively.

| | DSM15395 (PDB: 8DQM) | DSM2157 (PDB: 8DQN) |
|---|---|---|
| **Data collection** | | |
| Space group | $P2_12_12_1$ | $P2_12_12_1$ |
| Cell dimensions | | |
| *a*, *b*, *c* (Å) | 62.2 154.6 197.9 | 153.5 163.7 170.4 |
| α, β, γ (°) | 90.0 90.0 90.0 | 90.0 90.0 90.0 |
| Resolution (Å) | 98.96-2.70 (2.78-2.70) | 118.34-1.80 (1.86-1.80) |
| $R_{merge}$ | 0.033 (0.134) | 0.100 (0.868) |
| $R_{pim}$ | 0.033 (0.134) | 0.028 (0.245) |
| $I / \sigma I$ | 8.60 (0.65) | 9.68 (0.56) |
| $CC_{1/2}$ | 0.999 (0.972) | 0.999 (0.895) |
| Completeness (%) | 99.9 (99.9) | 98.2 (97.5) |
| Redundancy | 11.2 (7.9) | 13.6 (13.2) |
| **Refinement** | | |
| Resolution (Å) | 98.96-2.70 | 85.23-1.80 |
| No. reflections | 53406 (5216) | 386931 (38129) |
| $R_{work}$ / $R_{free}$ | 0.244/0.268 | 0.171/0.189 |
| No. atoms | 8662 | 25936 |
| Protein | 8510 | 22779 |
| Ligand/ion | 4 | 104 |
| Solvent | 152 | 3053 |
| *B*-factors | | |
| Protein | 41.18 | 31.00 |
| Ligands | 25.38 | 56.08 |
| Clashscore | 3.09 | 2.58 |
| Molprobity score | 1.33 | 1.19 |
| R.M.S. deviations | | |
| Bond lengths (Å) | 0.013 | 0.014 |
| Bond angles (°) | 1.80 | 1.83 |

**Table S1.** Statistics for crystallography data collection and structure refinement

Accompanying separate files:

**Table S2.xlsx**: Raw data used to generate the data in Table 1.

**Table S3.xlsx**: Raw data used to generate Fig. 2A, 3A and S2.
